# Evaluation of Upconverting nanoparticles towards heart theranostics

**DOI:** 10.1101/721035

**Authors:** Kermorgant Marc, Ben Salem Jennifer, Santelli Julien, Calise Denis, Oster Anne-Cécile, Lairez Olivier, Coudret Christophe, Verelst Marc, Gales Céline, Senard Jean-Michel, Beaudry Francis, Pavy-Le Traon Anne, Roux Clément, Mauricot Robert, Dina N. Arvanitis

## Abstract

Restricted and controlled drug delivery to the heart remains a challenge giving frequent off-target effects as well as limited retention of drugs in the heart. There is a need to develop and optimize tools to allow for improved design of drug candidates for treatment of heart diseases. Over the last decade, novel drug platforms and nanomaterials were designed to confine bioactive materials to the heart. Yet, the research remains in its infancy, not only in the development of tools but also in the understanding of effects of these materials on cardiac function and tissue integrity. Upconverting nanoparticles are nanomaterials that recently accelerated interest in theranostic nanomedicine technologies. Their unique photophysical properties allow for sensitive *in vivo* imaging that can be combined with spatio-temporal control for targeted release of encapsulated drugs.

Here we synthesized upconverting NaYF_4_:Yb,Tm nanoparticles and show for the first time their innocuity in the heart, when injected in the myocardium or in the pericardial space in mice. Nanoparticle retention and upconversion in the cardiac region did not alter heart rate variability, nor cardiac function as determined over a 15-day time course ensuing the sole injection. Altogether, our nanoparticles show innocuity primarily in the pericardial region and can be safely used for controlled spatiotemporal drug delivery.

Our results support the use of upconverting nanoparticles as potential theranostics tools overcoming some of the key limitations associated with conventional experimental cardiology.

## Introduction

Cardiovascular diseases (CVD), an umbrella term for a number of different pathologies including arteriosclerosis, coronary artery disease, arrhythmia, hypertension and heart failure, are the leading cause of morbidity and mortality in the world(1). Heart failure is a major cause of late morbidity and mortality after myocardial infarction (reviewed in (2)). Therapeutic interventions to prevent heart failure are of considerable active research. For example, treatment of the infarcted heart through experimental strategies has involved the delivery of growth factors, cytokines, and drugs to the infarcted cardiac tissues (reviewed in (3–5)).

Principal drug delivery methods indirectly target the heart. However, despite the efficiency of the usual routes of administration, they often result in off-target effects as well as limited concentrations and low retention of the factors in the desired area. To circumvent the low cardiac bioavailability and systemic effects observed by peripheral administration routes, targeted drug delivery approaches are currently considered. For instance, direct injections into the cardiac muscle, the myocardium, have been used to localize drugs or stem cells to infarcted regions(6). Drug vectorization to the heart can also be achieved by injection into the pericardial space which has a large potential for localized drug delivery (7). Both approaches for injection, whether in the intramyocardial (intraMY) and intrapericardial (intraPE) space, are used in experimental and clinical settings and are highly adaptable to the study design and expected endpoints. To date, despite significant advances in the development of effective cardiac therapeutic agents and in drug delivery technologies, enduring drug availability in the heart remains a complex and pervasive conundrum in research and clinical settings. As such, there is a need to develop drug-delivery and repeat-release approaches to deliver and maintain therapeutic materials to cardiac tissue with minimal adverse effects.

The need for spatiotemporal control for targeted release of various drugs that can be coupled with the capacity for imaging has accelerated interest in theranostic technologies. Theranostics is the integration of imaging (diagnostics) and drug delivery (therapy) for application in personalized medicine(8). In recent years, lanthanide-doped upconverting nanoparticles (UCNPs) have emerged as suitable tools for use in theranostics because of multiple features associated to the lanthanide ions. In particular, their unique optical properties originating from their electron configuration of the 4f shell(9). Indeed, the various lanthanide ions typically present a great number of energy levels and their excited states are usually long lived(10). Due to these spectroscopic features, these rare earth ions are in contrast to other chromophores whereby they accept energy in the near-infrared range (NIR; 980 nm, when doped with Ytterbium as the sensitizer) and emit at lower wavelengths than the excitation. When the nanocrystals are doped with Thulium as the activator, the main emission is observed in the biologically favourable 800 nm range(10). The illumination in the NIR, and anti-Stokes luminescence(11) is used to reduce the auto-fluorescent background allowing for improved signal-to-noise ratio. The narrow emission bandwidth of UCNPs allows ease of multiplexed imaging, and very limited to inexistent photobleaching making them relevant for longterm and repetitive imaging(12). In addition, UCNPs permit deep tissue penetration, for instance reported to be between 1.6 cm and 3.2 cm with minimal signal-to-noise background(13,14) because of the excitation in the NIR that is within the optical transparency window(15,16).

Given their advantageous photophysical properties, UCNPs have been exploited in a variety of *in vivo*(17) as well as in *in vivo* situations, ranging from luminescence-based imaging(18); to anticancer drug delivery(19); and, to mouse behavior control via optogenetics(20). In the context of targeted drug delivery, in order to minimize the drawbacks of conventional drug delivery with more specific and efficient target delivery, UCNPs as nanocarriers are promising tools(21,22). Few recent examples of heart-specific studies employing various nanomaterials are revealing a promising role for nanoparticle use to unravel the complexity and help treat CVDs(23–25). While still in its infancy, there is an imperative need to optimize the use of nanoparticles on the heart; with particular attention to the long-term conservancy of cardiac tissue and cardiac function *in vivo*.

The aim of the present study was to determine the innocuity of UCNPs to the heart and then explore their potential for time and space-controlled drug delivery. NaYF_4_:Yb,Tm NIR-emitting UCNPs were synthesized and used to measure the differential accumulation and distribution of the nanoparticles over a 15-day course following either intraperitoneal, intraMY or intraPE injections. Indeed, the pericardium, which is a thin layer that separates the heart from the thoracic cavity and provides structural support while also having a substantial hemodynamic impact on the heart, was a focal point of this study. Notably, we examined heart rate (HR), as determined by real-time electrocardiogram (ECG) monitoring, during the excitation upconversion phase that was repeated throughout the entire time course, to identify whether the UCNPs and or the laser beam localized to the chest area, have an impact on cardiac rhythm.

## Materials and methods

### Materials

Hydrated rare earth chlorides (99.9%), octadecene and oleic acid were from Alfa Aesar (Karlsruhe, Germany). NH_4_F, and NaOH were from Sigma-Aldrich (Saint Quentin Fallavier, France). PEO_6000_-PAA_6500_ was from Polymer Source (Montreal, Canada). All Other solvents were from Sigma Aldrich and of HPLC grade.

### Preparation of NaYF_4_:Yb,Tm Upconverting Nanocrystals

Upconversion nanocrystals were synthesized by a well-established high temperature liquid-phase protocol(26). YCl_3_.6H_2_O (315 mg, 1.03 mmol), YbCl_3_.6H_2_O (174 mg, 0.45 mmol), TmCl_3_.6H_2_O (3 mg, 7.8 μmol), dissolved in 500 μL of water were added to 24.5 mL of octadecene and 4.5 mL of oleic acid. The suspension was heated to 160°C under argon using a heating mantle, maintained at 160°C for 1 h, which yielded a clear light-yellow solution. After cooling to room temperature, 10 mL of a MeOH solution containing NH_4_F (222 mg), and NaOH (150 mg) were added dropwise while stirring. The suspension was heated and maintained at 50°C under a continuous argon flow for 30 min. The temperature was increased and maintained for 1 h at 100°C to allow for the evaporation of the methanol. The flask was then stoppered and 3 vacuum/argon cycles were applied using a high-vacuum pump. The heating mantle was then gradually increased to 310°C (the temperature reached 300°C after 15 min, 310°C in 30 min) for a total of 90 min. After cooling to room temperature, the crude reaction mixture was poured into 8 mL of ethanol. The particles were centrifuged at 9000 g and rinsed twice with ethanol. The solvent was then removed in a vacuum oven at 80°C, yielding 324 mg of a white powder.

### Preparation of Polyethylene oxide-polyacrylic acid (PEO-PAA)-coated UCNPs

5 mg of the above described particles were dispersed in 3 mL of toluene together with 5 mg of PEO_6000_-PAA_6500_. 3 drops of 1 M NaOH were added to help with polymer dissolution. This solution was transferred to a sealed vial and heated to 100°C for 1 h using a monomode microwave oven (monowave 300 Anton Paar, Les Ulis, France). Deionized water (1 mL) was then added and the aqueous phase was collected. Purification was achieved by three centrifugation steps at 9000 g. Following the last centrifugation, the particles were suspended in 2 mL of deionized water.

### Characterization of NaYF_4_:Yb,Tm Upconverting Nanocrystals

The particles were characterized by TEM microscopy (HT 7700 Hitachi), operating at 80 kV and 12 μA on standard formvar coated TEM grids (Electron Microscopy Sciences). Luminescence spectra were acquired on a modified PTI fluorimeter, where the traditional lamp has been replaced with a 980 nm CW laser (MDL-H-980, CNI lasers).

### Animals

Wild-type, 9-11week old males C57BL/6J (Envigo) were used. Animals were housed with food and water available ad libitum under standard 12h light/dark cycles. Animal procedures were approved by the national Animal Care and Ethics Committee (CE2A122 protocol number 2017092913349468) following Directive 2010/63/EU.

### Injections

For intraperitoneal injections, animals where briefly anaesthetized by a gas induction by 2% of isoflurane inhalant mixed with 1 L.min^−1^ 100% O_2_ and injected with 5 μL of a 1 μg.μL^−1^ solution of nanoparticles, and allowed to regain consciousness in their respective cages. For cardiac injections, anesthesia was induced using ketamine / xylazine (125/5 mg.kg^−1^) by intraperitoneal injection and maintained with 2% of isoflurane inhalant mixed with 1 L.min^−1^ 100% O_2_. The analgesic, buprenorphine (100 μg.kg^−1^), was administered subcutaneously. Using an operating microscope Zeiss OPM1 FC, tracheal intubation was performed for ventilation with the mini-wind (Harvard Apparatus). An incision of the 4^th^ intercostal space was performed to provide adequate exposure of the thoracic cavity. The left atrioventricular block was exposed and for intraMY injections a catheter was used to inject 5 μL of the nanoparticle solution into the myocardial wall of the apex of the left ventricle. For intraPE injections, a catheter was gently introduced within the pericardium and 5 μL of nanoparticle solution was injected. Evans Blue was used as a tracking dye to ensure lack of spillage. Controls consisted of phosphate buffer saline (PBS) injections into the respective areas. The intercostal space and skin surface were successively sutured using an Ethilon 6/0 thread (Ethnicon).

### BioImaging and Image processing

The setup consisted of a dark chamber supplied with an arrival for a precisely blended mixture of oxygen and isoflurane anesthetics. The compartment was also equipped with the Small Animal Physiological Monitoring system (Harvard Apparatus), which permitted rectal temperature control of the animal and continuous ECG readings. For upconversion and imaging, the chamber was fitted with a CCD camera (iKon-M 934, Andor) that was paired with an optical fiber and a beam expander (BE05M-B, Thorlabs) allowing for a 2.8 cm circular illumination area for excitation centered at 980 nm. All imaging sessions were performed with an excitation laser beam power density of 290 mW.cm^−2^ (controlled with a gentec power detector (PM130D, Thorlabs)) with 5 or 10 sec exposure times and were collected in control room temperature at 21°C. Hearts, including the pericardium, were microdissected, rinsed quickly in PBS and imaged with an excitation laser beam power density of 290 mW.cm^−2^.

A multicolor LUT was applied to the upconversion luminescence images, using a constant display range across all images. This upconversion image was then overlaid with an image recorded with white light illumination.

### Transthoracic Echocardiography

Mice were anesthetized by 2% of isoflurane inhalant mixed with 1 L/min 100% O_2_ to maintain light sedation throughout the procedure. They were immobilized ventral side up on a heating platform to maintain the body temperature at 37°C ± 0.5°C. Mice chests were shaved and warmed ultrasound gel was applied to the area of interest. Transthoracic echocardiography was performed using a Vevo 2100 system (VisualSonics) with a 40-MHz transducer by a trained user. Images were captured on cine loops at the time of the study and afterward measurements were done off-line. Cardiac ventricular dimensions were measured in M-mode and B-mode images 4 times for each animal. Left ventricle ejection fraction (LVEF) was calculated using parameters automatically computed by the Vevo 2100 standard measurement package. Measurements were obtained by an examiner blinded to the treatment of the animals. All measurements were performed excluding the respiration peaks and obtained in triplicate; mean values were used for data analyses.

### Heart rate variability (HRV) and power spectral analysis

For the acquisition of ECG, mice were anesthetized by inhalation of isoflurane at concentrations of 3% during the induction phase. Anesthesia was maintained at 1.5% and continuous recordings were obtained consisting of a minimum 10 min basal state, followed by laser usage (5 sec) and 10 min recordings using the Small Animal Physiological Monitoring system (Harvard Apparatus). ECG signal was obtained with three limb leads. The ECG signals were digitalized at 4 kHz, processed and monitored (Labchart v7, AD Instruments). The R-R interval was acquired from the ECG signal. Sections of stable HR (5 min before and after laser induction), free of noise and artifacts, were analyzed. HRV was assessed both in time and frequency domains.

Time domain measurements included the following metrics: the standard deviations of the R-R intervals (SDNN) and the root-mean square differences of successive R-R intervals (RMSSD). The SDNN corresponds to all the cyclic components responsible for the overall variability. RMSSD provides information about high frequency variations in HR. Overall, all these metrics reflect the autonomic status (Task Force, 1996).

Frequency domain analysis was performed using fast Fourier transform with 1,024 spectral points’ series using Welch’s periodogram with 50% overlapping window. Frequency domain analysis was analyzed in two separate spectral components: low frequency (LF: 0.15-1.50 Hz) and high frequency (HF: 1.50-5.00 Hz) bandwidths. Very low frequency (VLF: < 0.15 Hz) was excluded from the analysis. These spectral components were expressed in absolute values of power (ms^2^) and in normalized units (n.u.). The LF spectrum and the LF (n.u.) [(100 * LF power / (total power – VLF power)] have been considered as an index of cardiac sympathetic tone whereas HF spectrum and HF (n.u.) [(100 * HF power / (total power – VLF power)] reflected cardiac parasympathetic tone. The LF/HF value was obtained from ratio LF (ms^2^) / HF (ms^2^) and emphasized the sympathovagal balance(27).

### Analysis of cardiac tissue necrosis

After 15 days, mice were euthanized by cervical dislocation following isoflurane exposure and hearts were dissected. Hearts were quickly rinsed in cold PBS, and imaged by photo-upconversion to locate nanoparticles. To assess necrosis, cross sections were obtained using a Zivic Mouse Heart Matrix (Zivic) and 200 mm slices were incubated in 1% triphenyltetrazolium (TTC) at 37°C incubator for 10 min. Slices were gently removed from TTC and placed in 4% paraformaldehyde (Sigma) at 4°C for 24 h. The sections were rinsed gently in saline, placed within clear plastic sheets and images of TTC stained sections were captured using a digital scanner. Both sides were scanned and the digital photomicrographs were analyzed for white (damaged/necrotic) versus red (live) tissue using ImageJ, quantified and expressed as a percentage of the sum of necrotic areas from all sections by the sum of all areas from all sections multiplied by 100(28).

### Statistical analysis

To compare results between the injection groups, 2-way ANOVA was used. Paired t-test was used for within group comparison. Pearson’s correlation was used. A p-value < 0.05 was considered significant.

## Results

### Synthesis of the upconverting nanoparticles

Upconverting nanoparticles were synthesized using a well-established high-temperature wet chemical synthesis(26). Figure 1 shows the characterization of their main features. TEM (Fig. 1(a)) was used to establish that the synthesized particles are highly homogeneous in size and have slightly elongated oval shape, with dimensions of 34 nm in minor axis, 38 nm in major axis (Fig. 1(b)). Further insight of the crystal phase was obtained by powder X-ray diffraction and an exact match with hexagonal β-phase NaYF_4_ reference data was observed (Fig. 1(c)). The upconversion spectrum was recorded following 980 nm excitation (Fig. 1(d)). Beside a minor emission in the blue range (475 nm, ^1^G_4_→^3^H_6_ transition), and two in the red range (647 nm, ^1^G_4_→^3^F_4_; 695 nm, ^3^F_3_→^3^H_6_) the emission spectrum is dominated by a very strong emission centered at 800 nm (^3^H_4_→^3^H_6_ transition) which is typical of particles doped with thulium as emitter.

**Fig 1:**
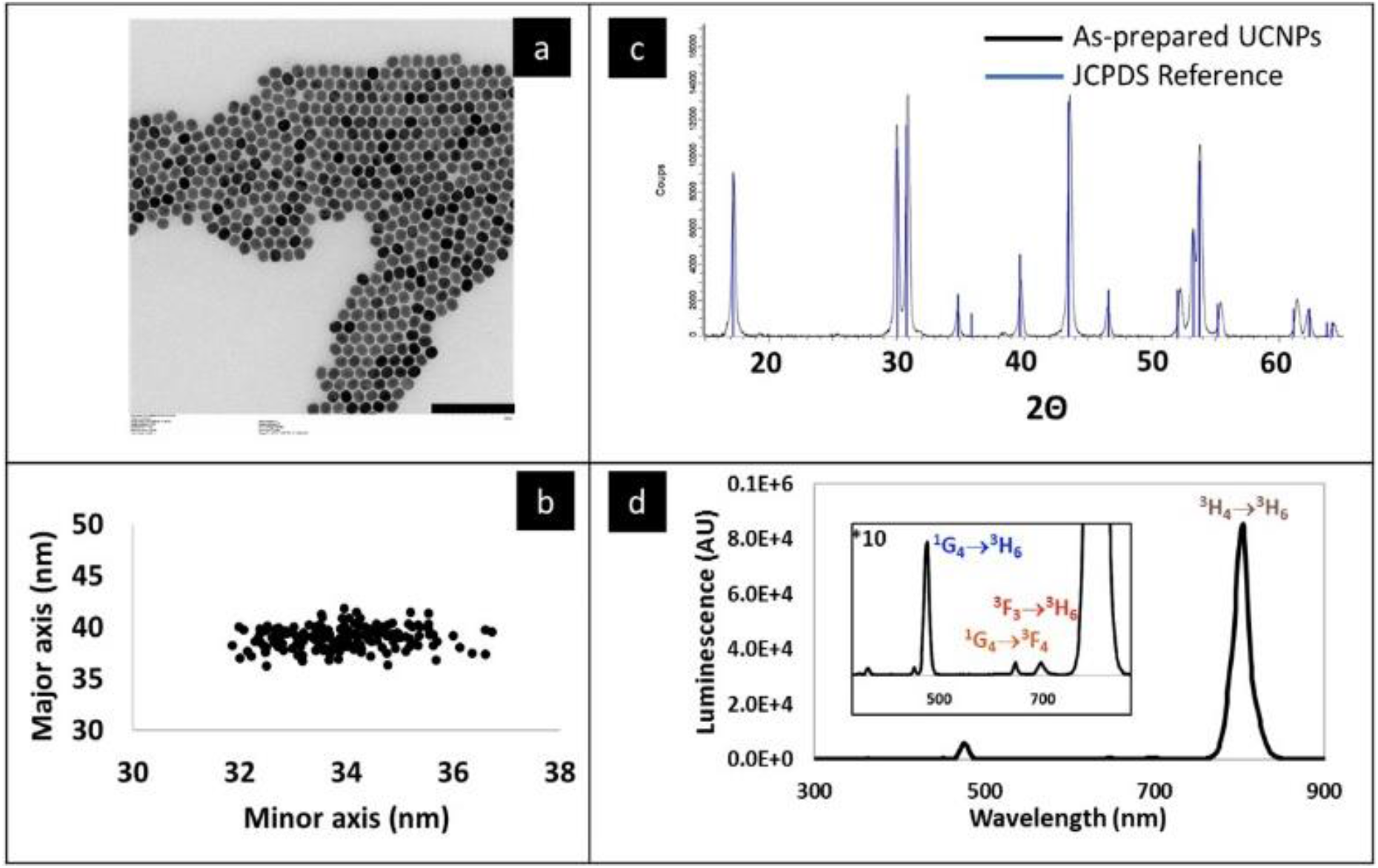
Characterization of the upconverting nanoparticles used in this study. A: Transmission electron micrograph. Scale bar = 200 nm. B: Individual sizes of the nanoparticles, along the minor and major axes. C: Powder X-ray diffractogram, and corresponding JCPDS reference (JCPDS No. 28-1192) data for NaYF_4_. D: Upconversion fluorescence spectrum of the particles, when excited by a 980 nm CW laser.

The particles initially covered in oleic acid ligands were rendered hydrophilic and biocompatible using a hydrophilic polymer coating, by the direct interaction of the OA-capped UCNPs with the macromolecules at 100°C (Polyethylene oxide-b-polyacrylic acid, PEO-PAA)(29).

### Experimental set-up

In this study, we investigated the fate of hydrophilic upconverting nanoparticles, following injection in three sites in mice: the peritoneum, the myocardium and the pericardium. The study is conceptually described in Fig. 2. In order to track nanoparticles, we assembled a bioimager that comprised a 980 nm laser source for excitation, which was passed through a beam expander in order to achieve an irradiated area of approximately 2.8 cm in diameter. The animal was anaesthetized using isoflurane and positioned according to the region of the body being examined. A physiological monitoring apparatus was implemented in the imager, which allowed for ECG to be recorded simultaneously to laser irradiation, and a rectal probe allowed for temperature control. The upconverted light was collected with a highly sensitive CCD camera, with peak quantum efficiency centered on 800 nm.

**Fig 2:**
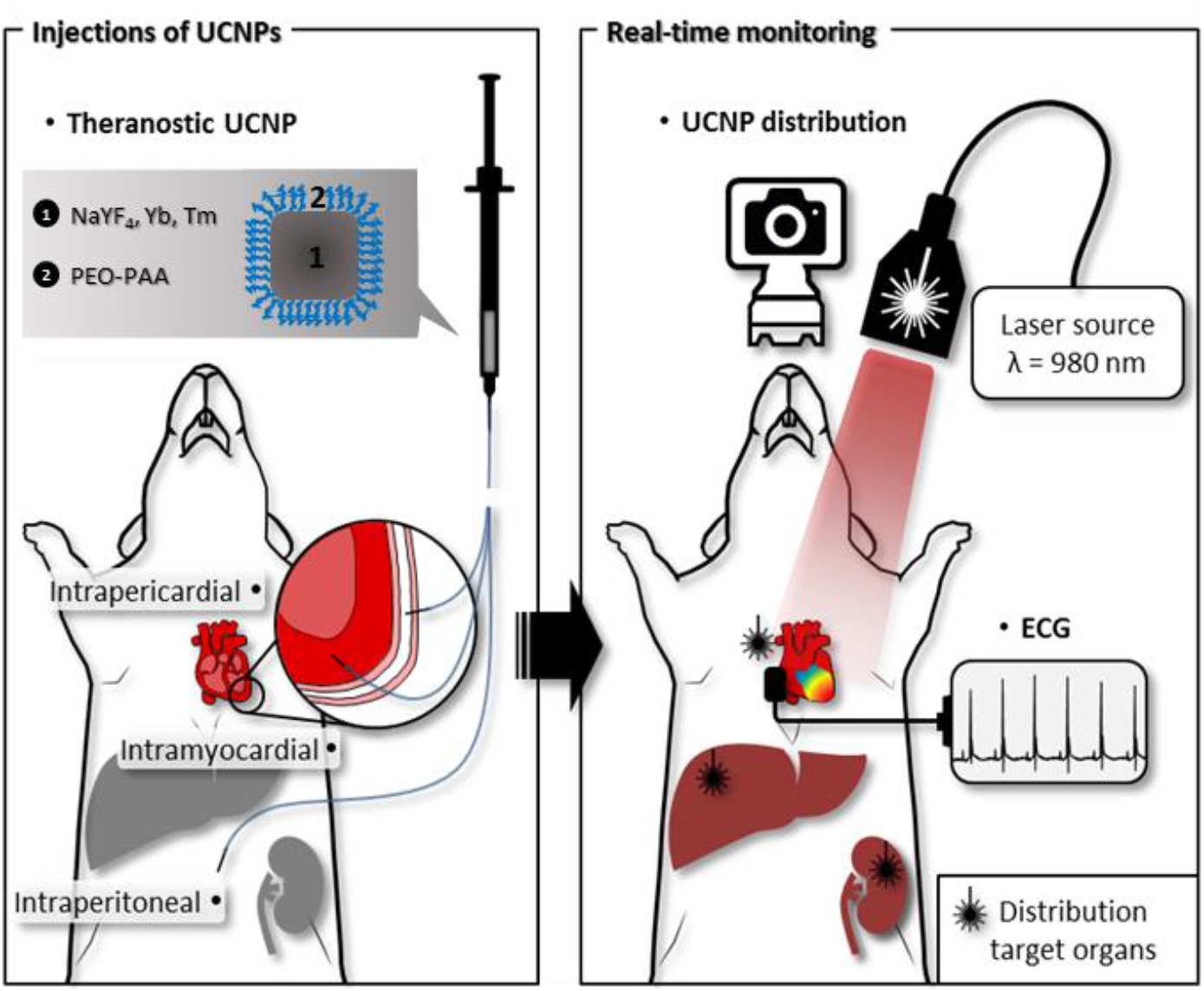
Schematic representation of the study design. UCNPs were injected in the myocardium, pericardium or peritoneum. Upconversion-based imaging was performed to image the UCNPs in the cardiac regions over a 15-day period. Retention and distribution of the UCNPs was tracked by upconversion in the heart, liver and kidneys. Continuous ECG monitoring was performed to determine the influence of the laser induced thermal effect on HR.

### Persistence of nanoparticles in the heart

Evans blue dye and bioimaging by upconversion immediately following the injection of the PEO-PAA coated nanoparticles were used to control for targeted placement and for any residual spillage of the injected solutions from the injection site. No non-specific leaks were detected in our experimental groups receiving intraMY or intraPE injections immediately following surgery. Time-course emission intensity was performed in the heart to determine the persistence of the nanoparticles within this region. Clearance of the nanoparticles was determined based on emission intensity at two regions: the kidney and liver. Following the injection by intraperitoneal route, which was included as control, we were unable to detect the presence of the nanoparticles in the heart at any of the time points measured. By 7-days post-injection there was no trace of nanoparticles in the subjects (Fig. 3(a)). Injection in the myocardium permitted the retention of the nanoparticles within the heart for 15 days, however, reduction of the upconversion luminescent emission at the heart was observed in the first 7 days post injection. While nanoparticles were retained in the heart muscle in all animals receiving intraMY injections, the dispersal of the UCNPs in these subjects were heterogeneously localized over time. Figure 3(b) is a representative animal, showing that nanoparticles persist in the intraMY space for 7 days, but are cleared from the heart within 15 days. IntraPE injections demonstrated retention in the heart region for 2 weeks with a slow clearance (Fig. 3(c)). Altogether, these data demonstrate that long-term nanoparticle retention in the heart is possible after intraMY and intraPE administration, but is optimally conserved in the cardiac region when localized to the pericardial space.

**Fig 3:**
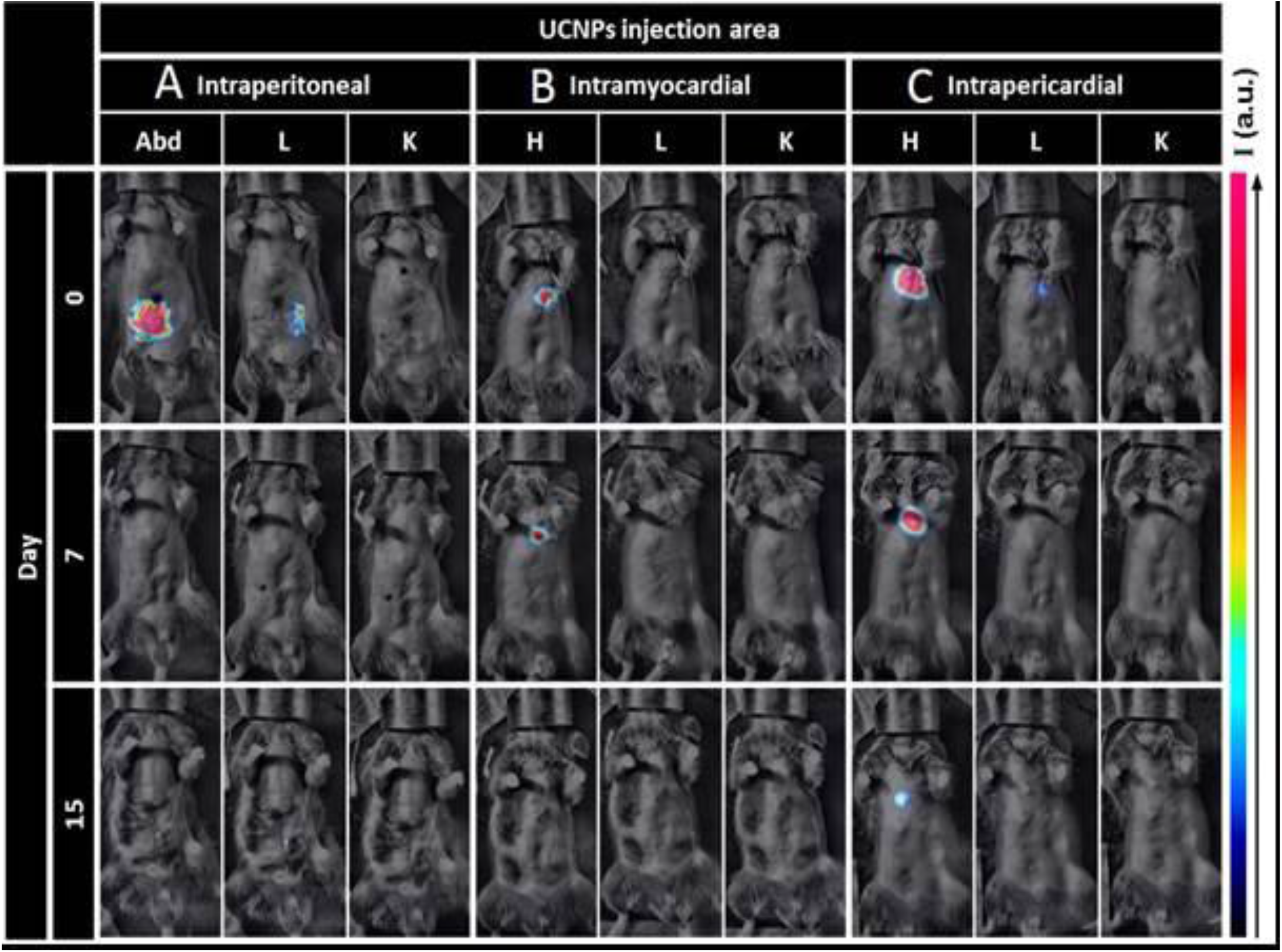
Spatio-temporal evolution of upconversion intensity. Representative images of the retention of UCNPs in the cardiac region and their biodistribution on day 0, day 7, day 15.

### Effects on body weight, Cardiac Function and Morphometry

Body weight was monitored throughout the experiment. As shown in Fig. 4(a), the mouse body weight of the control and experimental groups were similar showing a good macroscopic tolerance for the UCNPs. To further evaluate the *in vivo* effects of cardiac targeted nanoparticles on cardiac functions we implemented the Small Animal Physiological Monitoring System (Harvard Apparatus) along with the imaging setup. Real-time ECG monitoring during laser induced upconversion showed no effect on the heart rate in all subjects (Fig. 4(b)) even in the presence of UCNPs in the myocardium or pericardium (as confirmed in Fig. 3). No discernable variation was observed over the course of the 2 weeks post-injection following repeated upconversion on the same subjects (Fig. 4(c)). Therefore, we noted that presence of the UCNPs in addition to a brief (5 seconds) exposure of the laser beam power density of 290 mW.cm^−2^ to the chest region did not alter the ECG profile in UCNP-treated animals or controls.

**Figure 4.**
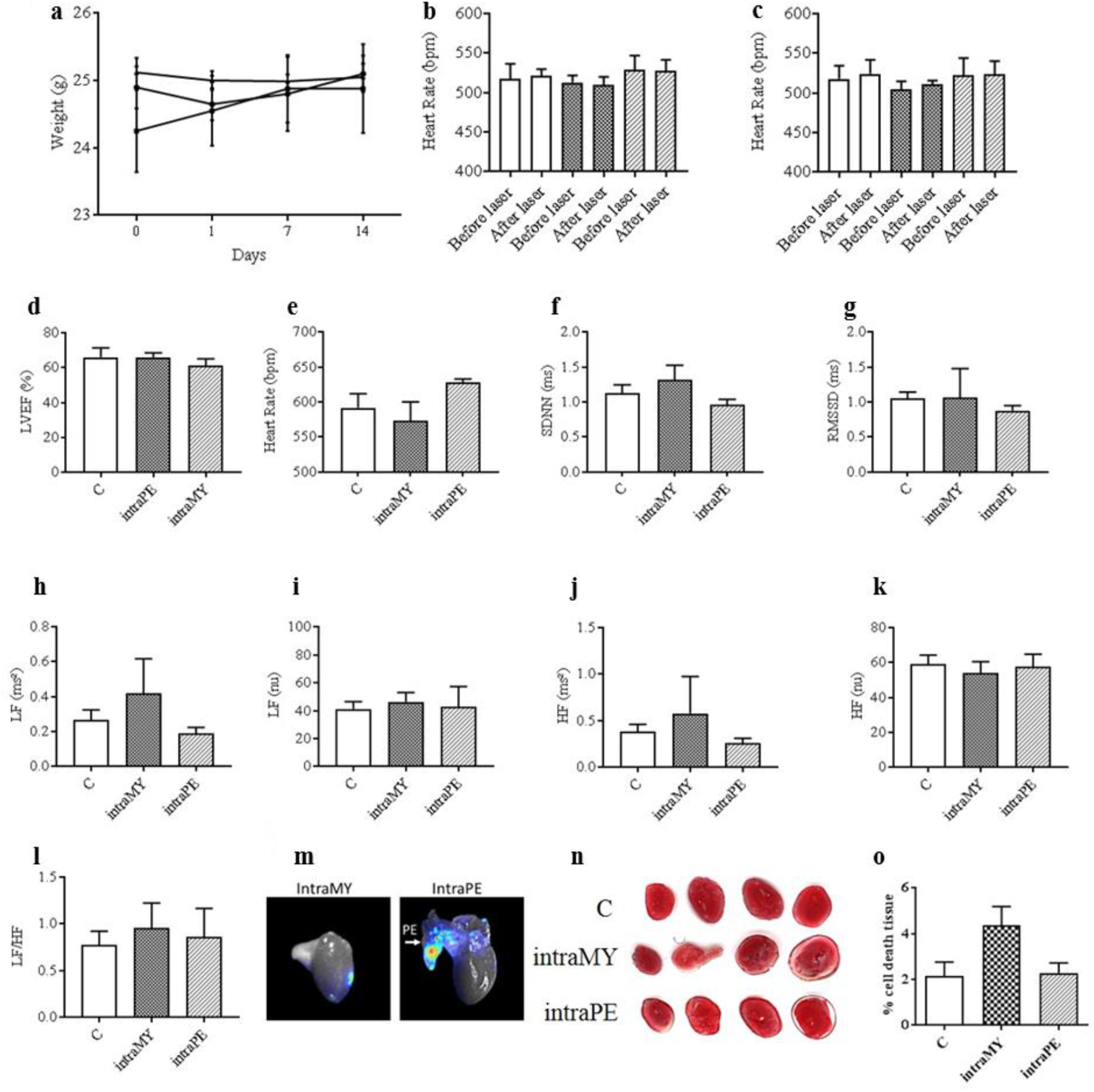
Cardiac cardiosympathovagal related parameters in nanoparticle treated animals. A Bodyweight changes in mice over a 14-day course, each point represents the mean ± SEM. B Percent left ventricle ejection fraction (LVEF) measured by echocardiography of control (C), intramyocardial (intraMY) and intrapericardial (intraPE) injected UCNPs. C HR measurements during laser induced upconversion on day 1 in C, and UCNPs targeted to the intraMY or IntraPE space. D HR measurements during laser induced upconversion on day 15 in C, and UCNPs targeted to the intraMY or IntraPE space. E End point HR measurements in C, and UCNPs targeted to the intraMY or IntraPE space. F, G SDNN and RMSSD analyses at day 15 post injection in C, intraMY and intraPE groups. H, I, J, K Low frequency spectrum, low frequency normalized units, high frequency spectrum and high frequency normalized units at day 15 post injection in C, intraMY and intraPE groups. L LF/HF ratio for all experimental groups. M isolated hearts showing that nanoparticles were localized to the myocardial injection site for intraMY administration, while the nanoparticles remained diffuse and associated with the pericardium and did not bind to the heart tissue when administered in the pericardial sac. N Macroscopic examination of cardiac tissue necrosis at 15 days post injection of the nanoparticles in C and experimental groups are shown. O Quantification of percent-tissue death.

Concerning *in vivo* endpoints, transthoracic echocardiography showed no effect of the interventions on the LVEF, a measure of LV muscle function, between the groups (Fig. 4(d)). ECG analyses showed no significant difference between experimental groups and controls in HR (Fig. 4(e)) or in any time domain measurements of HRV, either for SDNN (Fig. 4(f)) or RMSSD (Fig. 4(g)). Moreover, no significant change was noted in frequency domain measurements of HRV, including total power, LF (Fig. 4(h)), LF (n.u.) (Fig. 4(i)), HF (Fig. 4(j)), and HF (n.u.) (Fig. 4(k)), therefore, the sympathovagal balance (LF/HF ratio) was unmodified in the presence of nanoparticles (Fig. 4(l)). In an attempt to sublocalise the UCNPs within the heart following intraMY and intraPE injections, relevant hearts were dissected and the pericardia were reclined as shown in figure 4(m). It was observed that intraMY led to a localization of the UCNPs in the heart (Fig. 4(m), left), while UCNPs injected intraPE remained within the pericardium and did not bind to the heart itself (Fig. 4(m), right). Macroscopic examination of necrosis of cardiac tissues at 15 days post injection of the nanoparticles and quantification of percent-tissue death are respectively shown in Fig. 4(n) and 4(o) and evidence the absence of necrosis of cardiac tissue following pericardial sac injections, even if some tissue damage is perceivable in animals receiving intraMY.

## Discussion

We have synthesized NaYF_4_:Yb,Tm upconverting nanocrystals with an average size of 38 nm that exhibit spectrally sharp, large anti Stokes-shifted emission. For the persistence study, the particles, which are initially covered in oleic acid ligands, were rendered hydrophilic by coating with a layer of PEO-PAA copolymer. This double hydrophilic copolymer was anchored to the nanoparticles by multiple chelation from the polyacrylate segment. The second block, composed of PEO, ensured aqueous dispensability, and biocompatibility(30).

In an effort to identify the potential application of these UCNPs for heart specific photodynamic imaging, we decided to target the nanoparticles to the myocardium and/or the pericardial space. First, we assessed the potential of the nanoparticles to be maintained within the cardiac region. UCNPs injected in the myocardium exhibited release from the heart, yet there was no detectable accumulation in the liver or kidney. While intraMY injections permit highly localized delivery for clinical and research applications, incomplete retention, and even immediate leakage, from the heart have been reported for intraMY administration(31–33). This phenotype is attributed to the vascularization and dynamic environment of the myocardium. Improved, consistent and longer-term retention was observed for nanoparticles injected in the pericardial space. We found that nanoparticles injected into the pericardial sac result in highest retention. Pericardial sac injections are considered as a minimally invasive method to optimally localize therapeutics to the heart(7,34). UCNPs were retained in the pericardial region, and evaded lymphatic clearance even though they measured ~ 38 nm (Fig. 2), which is smaller than the depicted circular fenestrations of the parietal pericardium (diameter up to 50 μm (35)) therefore rendering their potential for several new and exciting applications. Our findings are in agreement with a recent study that showed BODIPY-containing PLGA nanoparticles administered intrapericardially were retained for a half-life of roughly 7-days and showed no microscopic detrimental consequence to the heart(36). Moreover, in terms of nanoparticle clearance, our findings correspond to other studies which show that the biodistribution and clearance of most nanoparticles *in vivo* result in their accumulation in the liver or kidney(37–39).

Despite favorable localization of the nanoparticles to the heart via intraMY or pericardial space injections, their influence on cardiac rhythm *in vivo* and over time, is unknown. Here we performed simultaneous and continuous ECG recordings with time-space controlled repetitive upconversion to the cardiac region and showed no adverse effects on HR. The frequency of the cardiac cycle is reflected as HR and is one of the most important physiological parameters that gives correct assessment of heart function.(40) Overall, nanoparticles in the myocardium or pericardial space did not alter the cardiac rhythm as detected by ECG analyses. End point experiments included echography, to visualize and assess cardiac function via the calculation of the LVEF(41), HRV analysis to identify cardiac rhythm abnormalities, and, autonomic nervous system assessment(42). No difference between UCNP-injected and control subjects was noticed. Though a slight decrease in the LVEF was observed in mice receiving direct intraMY injections of UCNPs as compared to pericardial space injections, neither of the administration routes resulted in cardiac muscle necrosis. Altogether, these data show that UCNPs can be localized and maintained in the heart for 15 days and can be subject to repeated excitation and upconversion with no effects on cardiac functioning or tissue integrity.

## Conclusions

Altogether, we provide the first evidence for multifunctional UCNPs that showed strong upconversion luminescence under 980 nm excitation, retention in the cardiac regions over the course of 15 days, biocompatibility, and high signal-to-noise ratio *in vivo*. The lengthy exposure of the UCNPs in the heart, and repeated excitation had no discernable effect on cardiac function. Future works are now aimed at applying our UCNPs to improve and multiplex with other imaging techniques, such as cardiac MRI. Our strategy has the potential to optimize targeted delivery of materials to the heart for biomarker identification in order to stratify heart failure and help enhance therapeutic strategies.

## Acknowledgments

We acknowledge core support from Animal facility ANEXPLO, CREFRE US06 Rangueil. This work was funded by the CNRS (Défi IMAG’IN-AAP 2017). Jennifer Ben Salem was supported by the Fondation de France. Grant number RAF18002BBA awarded to Dina N Arvanitis. Julien Santelli was supported by grants from CHROMALYS, INSERM and the “Région Midi-Pyrénées”. ANR is acknowledged for funding project “BLINK”: ANR-15-CE09-0020.

We thank Claire Pibourret and Baptiste Amouroux for their technical assistance in nanoparticle synthesis.

